# Resting-state heartbeat-evoked potentials are associated with Kalman-derived cardiac prediction errors

**DOI:** 10.64898/2026.05.13.724845

**Authors:** Takahiro Hirao, Kokono Terada, Mitsuhiro Miyamae, Makiko Yamada

## Abstract

The heartbeat-evoked potential (HEP) reflects the cortical processing of cardiac afferent signals. However, it remains unclear whether trial-level interoceptive prediction errors can be quantified directly from spontaneous resting cardiac fluctuations and whether these model-derived errors are associated with HEP amplitude. Here, we applied a Kalman filter, implemented as a sequential Bayesian estimation procedure, to resting-state EEG and ECG recordings from 21 healthy adults to estimate trial-by-trial signed prediction errors in RR-intervals. Positive prediction errors reflected unexpected cardiac deceleration, whereas negative prediction errors reflected unexpected cardiac acceleration. Cluster-based permutation tests showed that unexpected cardiac acceleration was associated with greater fronto-centro-parietal HEP amplitude than unexpected deceleration in an early post-R-peak window, spanning FC1, CP1, Pz, CP2, Cz, C4 and FC2 from 215 to 250 ms. A Bayesian linear mixed-effects model further indicated a credible negative association between signed prediction error and HEP amplitude after controlling for respiratory phase and preceding RR interval. In a secondary connectivity analysis, unexpected acceleration was associated with stronger Cz-to-frontal beta-band phase synchrony during a later post-R-peak window from 250 to 500 ms. Exploratory individual-difference analyses suggested that neuroticism was negatively correlated with late frontal HEP amplitude during unexpected acceleration, but not during unexpected deceleration or when trials were pooled across conditions. These findings demonstrate that spontaneous cardiac fluctuations can be used to derive trial-level computational estimates of interoceptive prediction error and that these estimates are reflected in early HEP amplitude. They further suggest that the cortical processing of unexpected cardiac acceleration may be related to individual differences in affective personality traits.

## Introduction

The heartbeat-evoked potential (HEP) is an event-related potential obtained by averaging electroencephalographic (EEG) signals time-locked to the R-peak of the electrocardiogram (ECG) (Park and Blanke 2019; Schandry, Sparrer, and Weitkunat 1986). The HEP is thought to reflect the cortical processing of afferent signals transmitted from the heart to the brain, and has attracted growing attention as a neurophysiological index of interoceptive processing. HEP amplitude has been associated with a range of psychological and cognitive processes, including cardiac interoceptive accuracy (Pollatos and Schandry 2004), emotional arousal (Luft and Bhattacharya 2015), and perceptual awareness (Park et al. 2014). However, methodological practices vary considerably across studies — including choice of reference electrode, baseline interval, analysis time window, and artifact rejection procedure (Coll et al. 2021; Park and Blanke 2019) — and the consistency of findings has been questioned (Coll et al. 2021).

A theoretical framework that offers a unified interpretation of HEP amplitude is interoceptive predictive coding (Barrett and Simmons 2015; Seth and Friston 2016). According to this framework, the brain constructs a generative model of interoceptive sensations based on prior experience and uses it to generate predictions about incoming cardiac afferent signals. The discrepancy between predicted and actual signals constitutes a prediction error. Within this framework, HEP amplitude is interpreted as reflecting the cortical processing of the prediction error generated at each heartbeat (Petzschner et al. 2019). Empirical support for this interpretation has accumulated (Banellis and Cruse 2020; Gentsch et al. 2019; Petzschner et al. 2019); however, each of these studies employed experimental tasks in which prediction error was indirectly manipulated by directing attention or varying emotional context.

According to interoceptive predictive coding theory, interoceptive processing proceeds through the following stages. First, the visceromotor cortex — encompassing the ventromedial prefrontal cortex (vmPFC), anterior cingulate cortex (ACC), and anterior insula — generates predictions about incoming cardiac afferent signals based on a generative model derived from prior experience (Barrett and Simmons 2015; Seth and Friston 2016). Actual cardiac signals ascend from peripheral sensory receptors, including baroreceptors, via the vagus nerve, nucleus of the solitary tract (NTS), parabrachial nucleus (PB), and thalamus to the posterior insula, which serves as the primary interoceptive cortex (Craig 2003). It is proposed that the posterior insula compares this ascending input against the prior prediction to generate a prediction error (Barrett and Simmons 2015). This prediction error is then processed by a cortical regulatory network comprising the anterior insula and ACC, which acts to minimize prediction error; top-down factors such as attentional state and emotional context modulate this process. The generative model is subsequently updated to inform future predictions, and homeostasis is maintained via autonomic responses mediated by descending projections from the vmPFC and anterior insula through the hypothalamus and brainstem (Barrett and Simmons 2015; Seth and Friston 2016).

The HEP has been used as a measure that captures the cortical response to cardiac interoceptive processing of this kind. Two components corresponding to the processing stages described above have been identified (Gautier, Latinus, and Briend 2025). An early component, observed with a fronto-central scalp distribution approximately 100–250 ms after the R-peak, reflects primary cardiac signal integration and is consistently observed regardless of task demands (Gautier et al. 2025). A later component, observed with a centro-parietal distribution at 250–500 ms, reflects higher-order processing that is modulated by top-down factors such as attention and arousal (Coll et al. 2021; Gautier et al. 2025).

The modulation of the late HEP component by top-down factors suggests that individual differences in interoceptive processing may be reflected in this component. Individual differences in interoceptive accuracy are known to be associated with affective style and vulnerability to psychiatric disorders (Critchley and Garfinkel 2017; Khalsa et al. 2018), and trait anxiety and depressive tendencies have been reported to modulate HEP amplitude even in healthy adults (Braun et al. 2025; Ito et al. 2019). However, these studies have focused on specific symptom dimensions, and the relationship between broader personality traits and the trial-by-trial processing of cardiac prediction errors in healthy adults remains unclear.

Based on the interoceptive predictive coding framework (Barrett and Simmons 2015; Seth and Friston 2016), it should be possible to quantify trial-by-trial prediction errors directly from spontaneous cardiac fluctuations during rest and to examine their correspondence with HEP amplitude. To our knowledge, however, no study has yet done so. The present study therefore employed sequential Bayesian estimation (Kalman filter) to compute trial-level RR-interval prediction errors from resting-state EEG recordings in healthy adults. We aimed to determine (1) whether cardiac prediction error — particularly its directional sign (unexpected acceleration versus deceleration) — is associated with HEP amplitude, and (2) whether this association varies as a function of personality traits.

## Methods

### Participants

Twenty-three individuals (12 women, 11 men; mean age = 42.4 years, *SD* = 13.5, range = 19–58 years) were enrolled in this study. Two participants were excluded: one due to poor ECG signal quality with unclear R-peak detection, and one due to excessive EEG noise rendering the data unsuitable for analysis, leaving 21 participants (11 men, 10 women; mean age = 42.8 years, *SD* = 13.2, range = 19–58 years) for analyses. Twenty participants were right-handed and one was left-handed, as assessed by the Edinburgh Handedness Inventory (mean laterality quotient = 84.1, *SD* = 37.0; Oldfield 1971), had normal or corrected-to-normal vision, and reported no history of neurological or psychiatric disorders. Written informed consent was obtained from all participants prior to participation. The study was approved by the Ethics Committee of the National Institutes for Quantum Science and Technology (QST).

### Procedure

After providing written informed consent and completing a pre-session questionnaire battery, participants were fitted with an EEG cap, and CGX-system electrodes were attached to record ECG and respiration signals. Following equipment setup, participants stood comfortably with eyes open for a 2-minute resting-state recording block. This pre-rest recording constitutes the dataset analyzed in the present study. Participants subsequently completed baseline and intervention gait measurement blocks (not analyzed here), interspersed with additional physiological and questionnaire assessments.

Vertical deviations between the two traces represent Kalman-derived prediction error. Blue segments indicate heartbeats with large unpredicted cardiac deceleration (i.e., an RR interval substantially longer than predicted), and red segments indicate heartbeats with large unpredicted cardiac acceleration (i.e., an RR interval substantially shorter than predicted).

### Questionnaires

Five subscales from the Ten Item Personality Inventory (TIPI; Gosling, Rentfrow, and Swann, 2003) were administered: Extraversion, Agreeableness, Conscientiousness, Neuroticism, and Openness to Experience. These five traits were selected a priori as representing the standard Big Five personality dimensions. False Discovery Rate correction was applied across all 5 TIPI traits simultaneously within each analysis unit.

Two additional affective trait measures were included as exploratory variables to probe finer-grained aspects of positive psychological characteristics: the Future Prediction task scale (FPT), yielding a optimism thinking (Isato et al. 2026), and the Above Average Effect scale (AAE).

### Physiological recordings

EEG was recorded using a BrainVision LiveAmp amplifier (Brain Products, Germany) with BrainVision Recorder Professional software (v. 1.21.0402). Twenty-eight active dry electrodes were placed at standard 10–20 positions (Fp1, Fp2, F7, F3, Fz, F4, F8, FC5, FC1, FC2, FC6, T7, C3, Cz, C4, T8, CP5, CP1, CP2, CP6, P7, P3, Pz, P4, P8, O1, Oz, O2), referenced online to the right mastoid (M2). Three additional channels recorded vertical (vEOG) and horizontal (LHEOG, RHEOG) electro-oculogram. Signals were digitized at 500 Hz using a 24-bit ADC (voltage resolution: 0.041 µV/bit), with a hardware low-pass filter of 131 Hz. The ECG and respiratory signals were recorded simultaneously using a CGX system (256 Hz sampling rate) and synchronized with the EEG.

### Data analysis

#### EEG analysis

Processing for resting data was mainly performed using MNE-Python. Raw EEG was high-pass filtered at 0.1 Hz. Data were re-referenced to the average of all EEG electrodes. Continuous data segments containing peak-to-peak EEG amplitude exceeding 150 µV were annotated as bad. Muscle artifacts were first attenuated using Canonical Correlation Analysis (CCA; De Clercq et al., 2006), in which components were ranked by temporal autocorrelation and those with a spectral slope above −0.59 (indicative of muscle activity; Fitzgibbon et al. 2016) and autocorrelation below 0.5 were zeroed and reconstructed. A copy of the CCA-cleaned data was additionally bandpass filtered at 2–100 Hz for ICA fitting. ICA was performed using the extended Infomax algorithm. Components were classified using ICLabel (mne-icalabel; Pion-Tonachini, Kreutz-Delgado, and Makeig 2019); those with brain probability < 0.70 were submitted to targeted wavelet-enhanced ICA (wICA; Castellanos and Makarov, 2006), in which artifact-related wavelet coefficients were hard-thresholded and subtracted from the component time series before back-projection to scalp space. A low-pass filter at 30 Hz was applied to the cleaned data.

EEG data were epoched from −200 ms to +600 ms relative to each accepted R-peak (baseline: −130 to −30 ms). Epochs were rejected if peak-to-peak amplitude exceeded 75 µV at any of nine fronto-central and parieto-central electrodes (F3, Fz, F4, C3, Cz, C4, P3, Pz, P4), or if they overlapped with annotated bad segments. A surface Laplacian (Current Source Density; CSD) transform was applied using a spherical spline (µV/m^2^) to improve spatial specificity and reduce volume-conduction effects.

#### ECG and respiration

The ECG signal was bandpass-filtered at 5–35 Hz. R-peaks were detected using a custom script designed to detect all R-peaks excluding segments in which signal quality was insufficient for reliable R-peak identification. RR intervals together with concurrent respiratory phase were stored for use in all subsequent analyses.

#### Cardiac Predictor: Kalman Prediction Error

To operationalize interoceptive prediction error at the single-trial level, we employed a Bayesian state-space framework. Under interoceptive predictive coding theory, the brain maintains an internal generative model of cardiac dynamics from which predictions of upcoming afferent signals are derived; any mismatch between the predicted and actual heartbeat constitutes a prediction error whose cortical representation is reflected in HEP amplitude (Barrett and Simmons 2015; Seth and Friston 2016). A Kalman filter provides sequential, one-step-ahead Bayesian predictions that directly operationalize this construct, conditioning each predicted interval on the estimated autocorrelative structure of the preceding RR sequence.

RR intervals exhibit serial autocorrelation at the beat-to-beat timescale (Berntson et al., 1997; Task Force, 1996). To provide a parsimonious generative model for the Kalman filter, we adopted a first-order autoregressive model, which captures the dominant one-lag dependency in the RR sequence. The fitted first-order autoregressive coefficients were embedded in a scalar Kalman filter to generate one-step-ahead predictions:

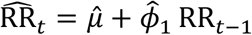

The prediction error at each beat was:

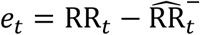

Positive values indicate unexpected deceleration; negative values indicate unexpected acceleration. Prediction errors were z-standardized within each participant and trials were split into unexpected deceleration (≥ median) and unexpected acceleration (< median) conditions.

### Statistical analysis

#### HEP Cluster-Based Permutation Test

To examine whether HEP amplitude differed between prediction error conditions, subject-level difference waveforms (unexpected deceleration − unexpected acceleration) were submitted to nonparametric spatio-temporal cluster permutation tests (1,000 permutations; one-sample t-test against zero). Spatial adjacency was defined as Euclidean distance ≤ 0.054 m among the 28 CSD channels. Consistent with Zaccaro et al. (2024), tests were conducted separately for early (100–350 ms) and late (350–600 ms) post-R-peak windows. The cluster-forming threshold was set at one-tailed p < 0.01, and clusters with a permutation p < 0.05 were considered significant. Positive-direction (deceleration > acceleration) and negative-direction (deceleration < acceleration) clusters were tested separately using one-tailed tests in each window.

#### Bayesian LMM on the Significant HEP Cluster

To quantify the trial-level relationship between Kalman-derived prediction error and HEP amplitude at the significant cluster while controlling for the influences of cardiac and respiratory covariates, a Bayesian linear mixed-effects model (LMM) was fitted to per-trial cluster-mean CSD amplitude. The model included Kalman-derived prediction error as the predictor of interest, respiratory phase and the preceding RR interval as covariates, and by-participant random intercepts and random slopes for Kalman-derived prediction error:

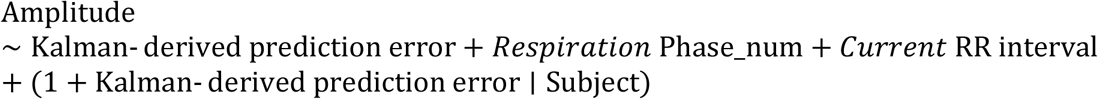

Posterior inference was performed using the No-U-Turn Sampler (NUTS; 2,000 warm-up iterations + 2,000 sampling draws; 4 chains; Bambi library). A fixed effect was considered credible if its 95% Highest Density Interval (HDI) excluded zero.

#### wPLI Connectivity Analysis — Cz Seed

To investigate whether the cortical processing of cardiac prediction error modulates large-scale oscillatory coupling, wPLI was computed using Cz as the connectivity seed — the channel corresponding to the peak-t statistic within the significant HEP cluster and thereby serving as the central locus of prediction error-related cortical activity. Epochs were split into unexpected deceleration and unexpected acceleration conditions using a within-subject median split of prediction error. Per-subject wPLI was estimated for 27 target channels across three frequency bands — theta (4–8 Hz), alpha (8–13 Hz), and beta (13–30 Hz) — and two non-overlapping post-R-peak time bins (early: 0–250 ms; late: 250–500 ms).

Condition-difference wPLI was computed as Δ wPLI = wPLI(unexpected deceleration) − wPLI(unexpected acceleration). Positive values indicate stronger Cz–target phase coupling during deceleration; negative values indicate stronger coupling during acceleration. Δ wPLI values were tested against zero using nonparametric spatial cluster permutation tests (1,000 permutations; spatial adjacency: Euclidean distance ≤ 0.054 m) conducted separately per band × bin combination. The cluster-forming threshold was set at one-tailed p < 0.01, and clusters with a permutation p < 0.05 were considered significant.

#### Condition-Split HEP Amplitude × Personality Trait

To examine whether individual differences in personality modulated condition-specific HEP amplitude, cluster-region mean CSD amplitude was extracted separately for the unexpected deceleration, unexpected acceleration, and deceleration − acceleration difference conditions. Each measure was correlated with the 5 TIPI subscales using Spearman’s ρ, with FDR-BH correction applied across all 5 traits within each cluster × condition combination.

To gain finer-grained insight into how the brain processes cardiac prediction error, analyses were additionally conducted at the level of individual electrodes within the significant connectivity clusters, separately for signed and absolute prediction error splits. As a further exploratory analysis targeting the processing of positive psychological traits, the Future Prediction Task scale (FPT) and the Above Average Effect scale (AAE) difference scores — measures reflecting positive-over-negative bias — were correlated with condition-split HEP amplitude using the same approach (see Supplementary document).

## Results

### HEP Cluster-Based Permutation Test: unexpected acceleration vs. deceleration

Kalman-derived prediction errors were computed from each participant’s observed RR-interval sequence and used to classify every trial as unexpected deceleration (prediction error ≥ within-participant median) or unexpected acceleration (prediction error < median; Figure 2). A cluster-based permutation test comparing HEP amplitude between these two conditions revealed one significant negative cluster in the early post-R-peak window (100–350 ms), spanning electrodes FC1, CP1, Pz, CP2, Cz, C4, and FC2 over the 215 to 250 ms interval (p_cluster = 0.014). The negative direction indicates that HEP amplitudes in this cluster were reduced when beats arrived later than predicted (unexpected deceleration) relative to beats arriving earlier than predicted (unexpected acceleration). No significant clusters were observed in the late window (350–600 ms) or for the positive-tail test (Figure 3).

**Figure 1.**
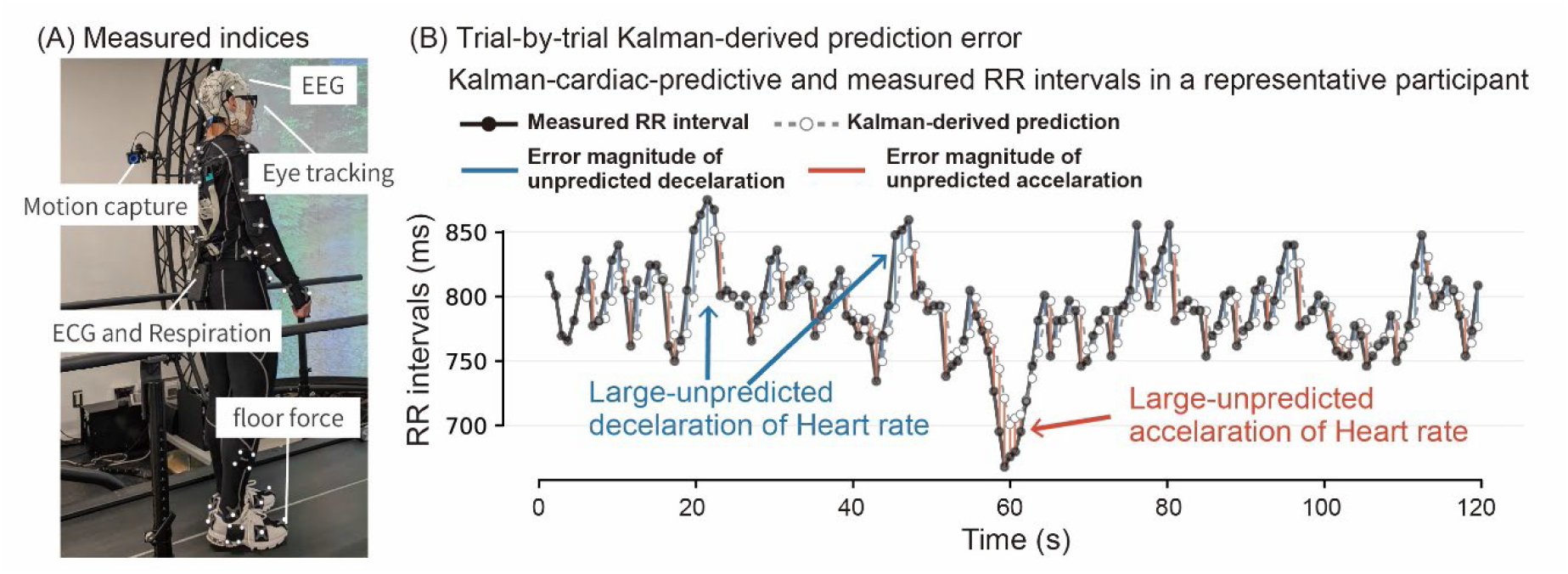
Experimental setup and trial-by-trial cardiac prediction error. (A) Multimodal measurement system used in a larger study. In the present study, only EEG and ECG/respiration recordings were analyzed. (B) Representative time series of measured RR intervals (black line with filled circles) and the Kalman filter-derived cardiac prediction (gray dashed line with open circles) for a single participant.

**Figure 2.**
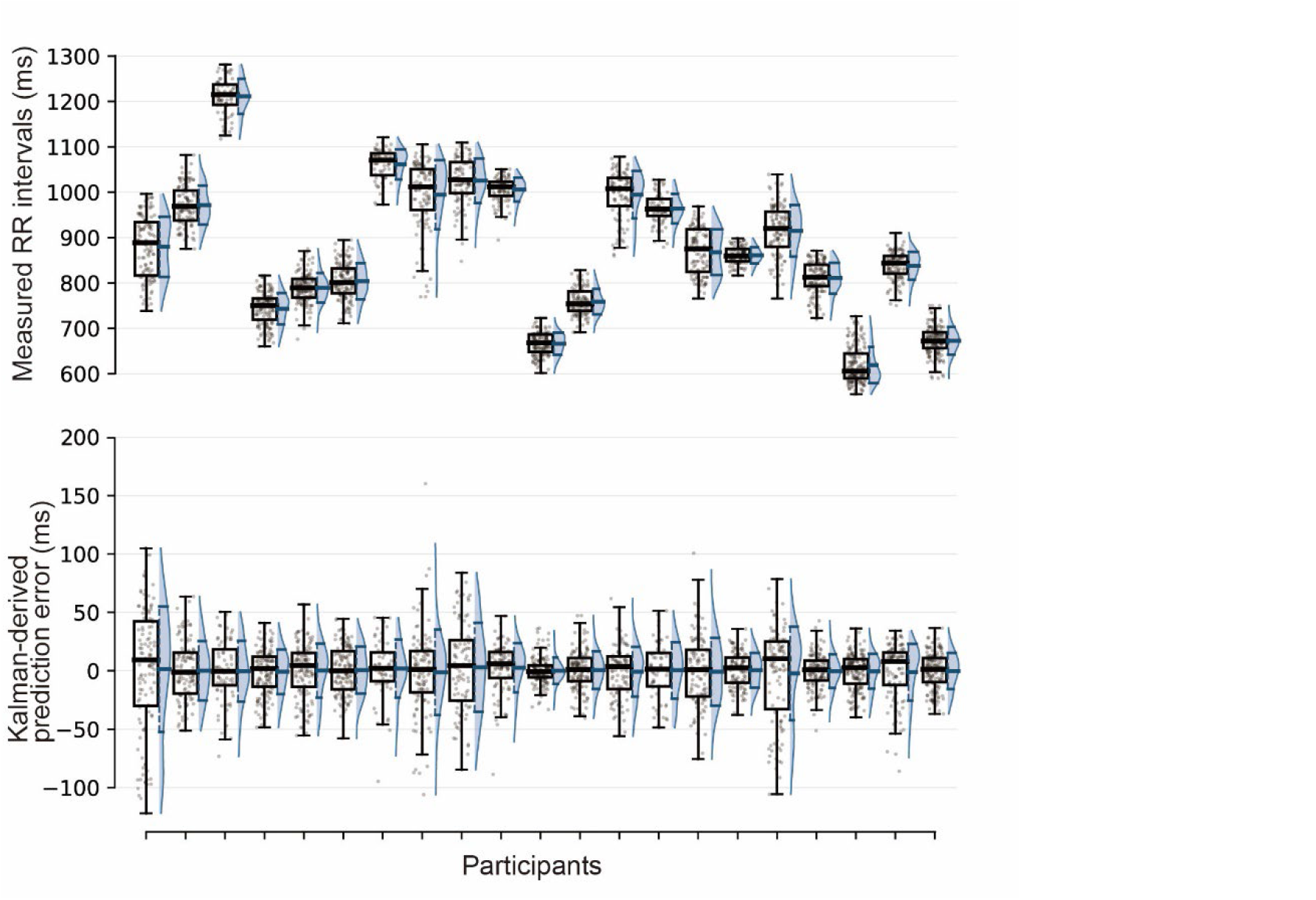
Per-participant trial-level distributions of measured RR intervals and Kalman-derived prediction error. Each panel shows per-trial data per participant; box plots indicate median and interquartile range; violins show kernel density estimates. The prediction error distribution was centered near zero within each participant, confirming balanced trial counts across conditions.

**Figure 3.**
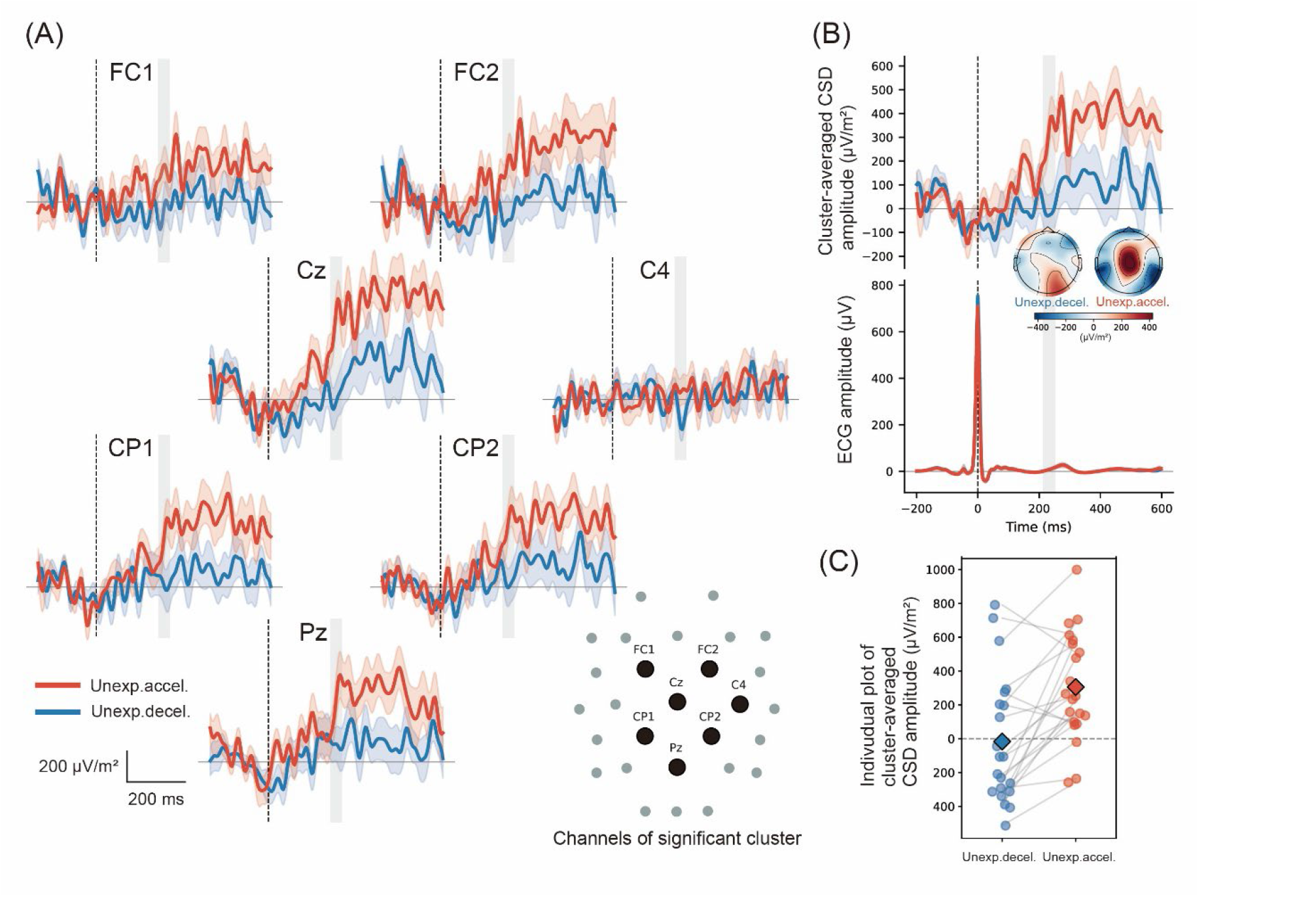
Cluster permutation test result for unexpected deceleration vs. unexpected acceleration. (A) Grand-average CSD waveforms (±1 SEM shading) and scalp topography of grand-average CSD are shown for each electrode in the cluster (215– 250 ms; FC1, CP1, Pz, CP2, Cz, C4, FC2; p_cluster = 0.014). (B) The cluster-mean waveform averaged across all significant electrodes; the bottom panel shows the grand-average ECG waveform per condition. (C) Individual plots for unexpected deceleration vs. unexpected acceleration. Each point represents a participant’s mean CSD amplitude (µV/m^2^) in the respective condition. Lines connect paired observations within participants. Diamonds indicate grand means.

### Bayesian LMM: Kalman prediction error Effect

Figure 4A illustrates per-participant trial-level distributions of HEP amplitude at the significant cluster. To quantify the trial-level effect of Kalman-derived prediction error on cluster amplitude while accounting for cardiac and respiratory covariates, a Bayesian LMM was fitted. A credible negative fixed effect of prediction error was observed (posterior mean = −6.54 µV/m^2^ per z-unit, 95% HDI = [−11.45, −2.18]; Figure 4B), indicating that each standard-deviation increase in prediction error (unexpected deceleration) was associated with a reduction of approximately 6.5 µV/m^2^ in cluster HEP amplitude in the 215 to 250 ms window.

**Figure 4.**
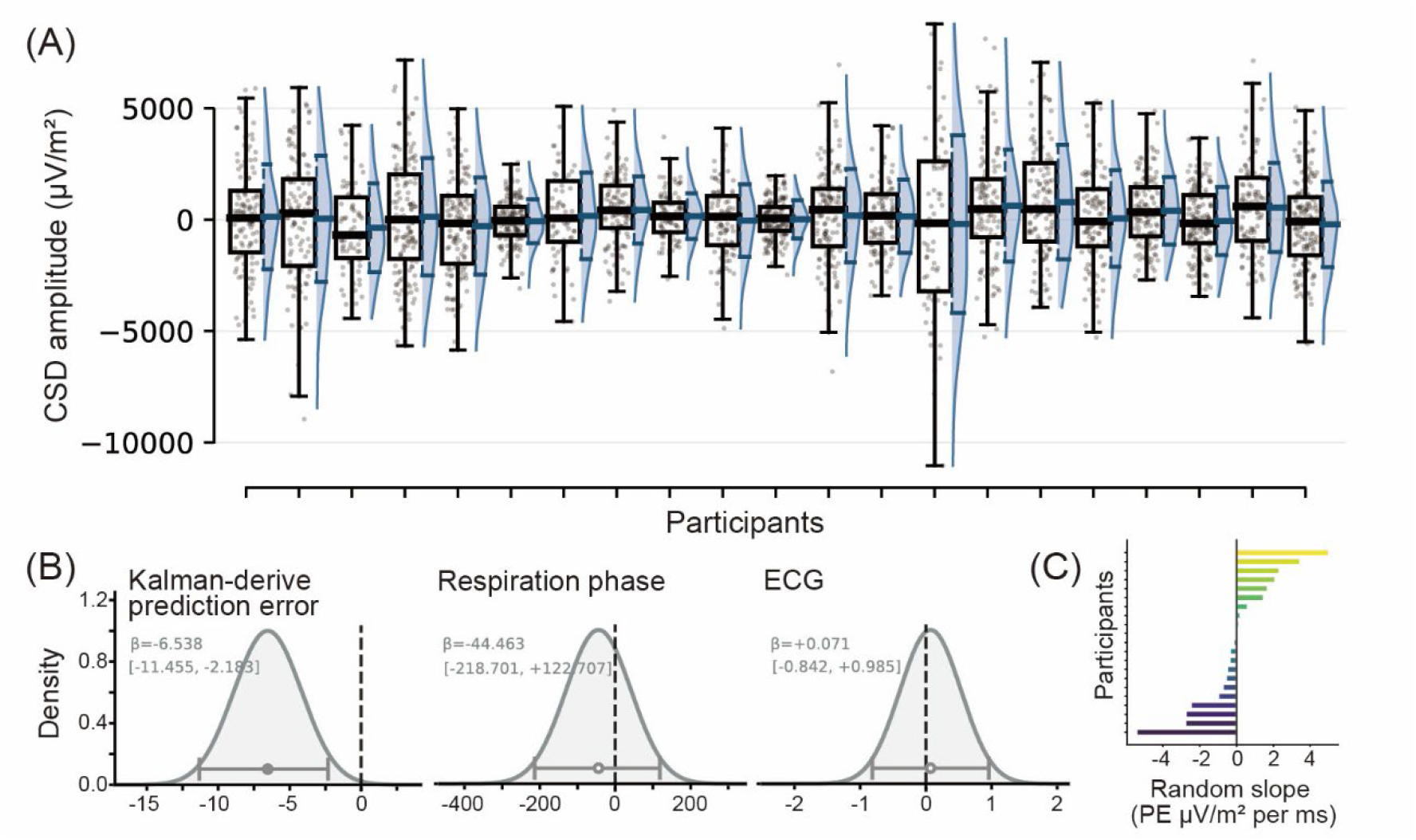
(A) Per-participant trial-level distributions of HEP amplitude at the significant cluster (215–250 ms; FC1, CP1, Pz, CP2, Cz, C4, FC2; µV/m^2^). Each panel shows per-trial data per participant; box plots indicate median and interquartile range; violins show kernel density estimates. (B) Posterior distribution of the fixed effect of the Bayesian linear mixed-effects model (LMM). The posterior mean (circle) and 95% equal-tailed interval (ETI; horizontal bar) are overlapped on the posterior density. (C) Per-subject random slopes for the Kalman-derived prediction error predictor in the Bayesian LMM. Each bar represents the posterior mean of the subject-specific deviation from the group-level fixed effect. Subjects are sorted by slope magnitude.

The Bayesian LMM also yielded per-subject random slopes for Kalman-derived prediction error, capturing each participant’s individual-level deviation from the group-average prediction error –amplitude relationship (range: −5.42 to +5.00 µV/m^2^ per z-unit; Figure 3C). To examine whether these individual differences in prediction-error sensitivity were associated with personality, per-subject random slopes were correlated with the five TIPI subscales; no association survived FDR correction (all p > 0.37).

### wPLI Condition-Difference Connectivity (Cz Seed)

Using Cz — the central electrode of the significant HEP cluster — as the connectivity seed, we examined whether prediction error-related beta-band coupling differed between unexpected deceleration and unexpected acceleration conditions. Two significant negative beta-band clusters were found in the late post-R-peak time bin (250–500 ms): a frontal cluster comprising electrodes Fz, FC2, and F4 (mean ΔwPLI = −0.063, p_cluster = 0.004), and a parietal-central singleton comprising electrode CP1 (mean ΔwPLI = −0.065, p_cluster = 0.046). No significant clusters were observed in the early time bin (0–250 ms) or in theta or alpha bands. The negative direction indicates stronger Cz-to-target beta phase coupling during unexpected acceleration than unexpected deceleration beats. Because CP1 is itself a constituent electrode of the significant HEP cluster centered on Cz, it was excluded from subsequent personality trait analyses to avoid potential circularity. Connectivity diagrams and cluster-region waveforms are shown in Figures 5A and B, respectively.

**Figure 5.**
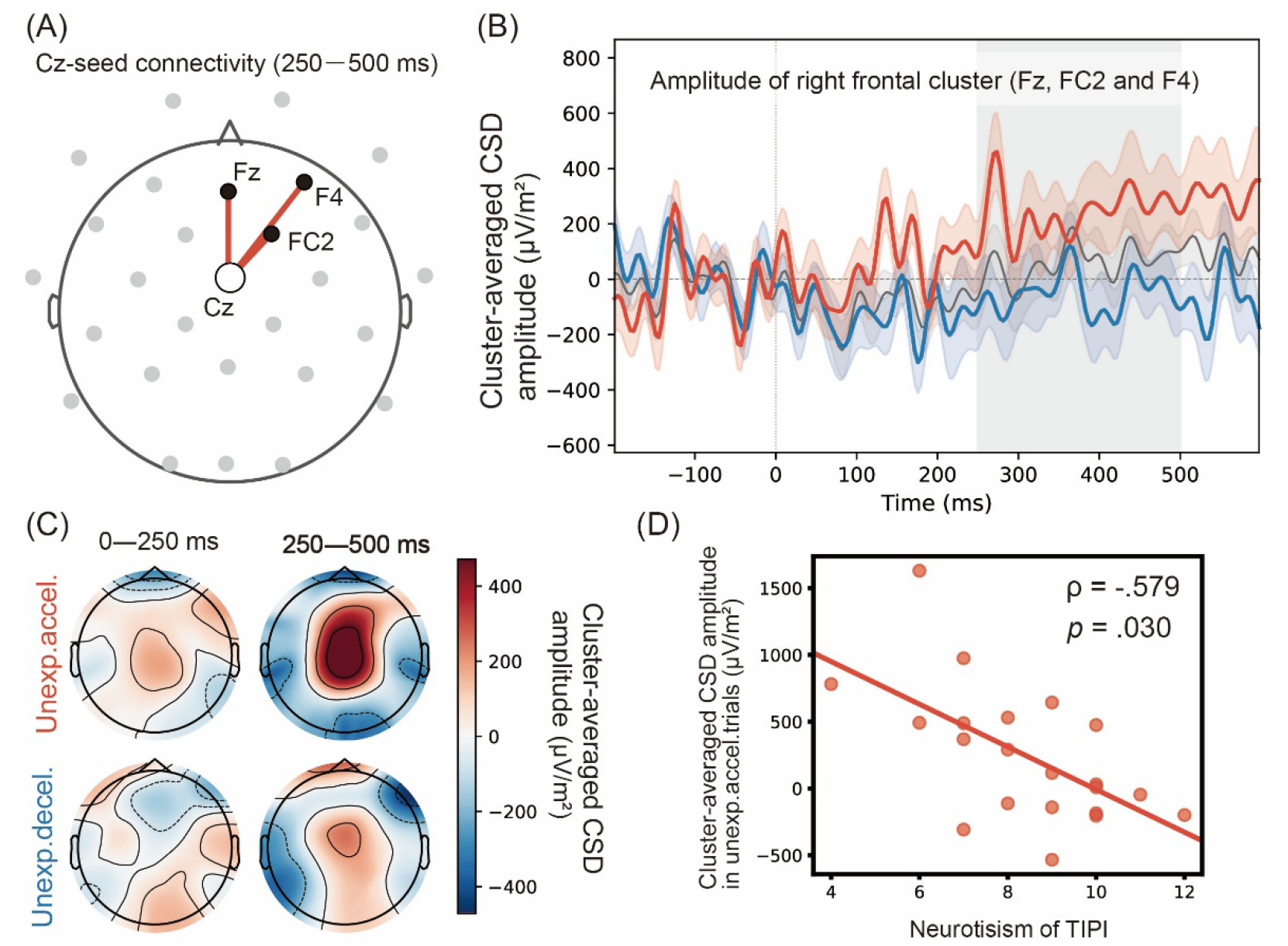
Results of connectivity. (A) Cz-seed wPLI connectivity diagram (beta band, 250–500 ms post-R-peak). Coloured edges indicate significant Δ wPLI clusters; edge thickness is proportional to mean |ΔwPLI|. (B) Cluster-region grand-average wPLI waveforms (unexpected deceleration vs. unexpected acceleration Kalman-derived prediction error; ±1 SEM shading). (C) Scalp topography of grand-average CSD HEP amplitude (µV/m^2^) for the unexpected deceleration and unexpected acceleration Kalman-derived prediction error conditions across two post-R-peak time bins (0–250 ms; 250–500 ms). (D) Spearman correlation between cluster-mean HEP amplitude at the frontal cluster with significant Cz-seed beta connectivity (Fz, FC2, F4; 250–500 ms post-R-peak) during the unexpected acceleration condition and Neuroticism (ρ = −0.579, p = 0.030). Each point represents one participant.

### Condition-Specific HEP Amplitude and Personality Traits

A significant negative association was observed between HEP amplitude during the unexpected acceleration condition and Neuroticism at the frontal beta cluster (Fz, FC2, F4; 250–500 ms): ρ = −0.579, p = 0.030 (Figure 5D). Participants with higher Neuroticism scores showed lower HEP amplitude at this cluster specifically during unexpected acceleration beats. This association was not significant for the unexpected deceleration condition (ρ = −0.219, p = 0.565) or the deceleration − acceleration difference condition (ρ = −0.337, p = 0.339), and did not reach significance when all trials were pooled regardless of prediction error level (ρ = −0.504, p = 0.099), indicating a condition-specific rather than general HEP–Neuroticism link.

As an exploratory finding, a significant negative association was observed between HEP amplitude at FC2 (250–500 ms) during high absolute prediction error beats and the Future Positive Task score (FPT; ρ = −0.455, p = 0.038; Supplementary Figure S1). No FDR-significant associations were observed for the Above Average Effect scale (AAE) (Figure S1).

## Discussion

The directional sign of the Kalman filter-derived prediction error was associated with HEP amplitude. During unexpected acceleration, HEP amplitude at the fronto-centro-parietal cluster (FC1, CP1, Pz, CP2, Cz, C4, FC2; 215–250 ms) was greater than during unexpected deceleration. In the Bayesian LMM, the association whereby greater unexpected deceleration was linked to lower HEP amplitude — consistent in direction with the cluster-based permutation test result — was maintained after controlling for respiratory phase and instantaneous heart rate (preceding RR interval; posterior mean = −6.54 µV/m^2^; 95% HDI [−11.45, −2.18]). This result raises the possibility that cortical processing of the prediction error occurred within the 215–250 ms window. Whereas prior studies demonstrated HEP modulation by indirectly manipulating prediction error through experimental tasks (Gentsch et al. 2019; Petzschner et al. 2019), the present study showed that prediction error computed directly from spontaneous resting-state cardiac fluctuations is associated with HEP amplitude. Given that 215–250 ms falls within the early component (∼100–250 ms) of the two-component model proposed by Gautier et al., (2025), the present findings suggested that directional discrimination of prediction errors derived from natural resting-state cardiac variability already emerges at the stage of primary cardiac signal integration. In other words, the early HEP component may not merely register incoming cardiac afferent signals at the cortical level, but may also reflect — at this initial stage — the mismatch between predictions about heartbeat timing and the actual arriving signal.

Unexpected acceleration and unexpected deceleration are thought to reflect distinct branches of the autonomic nervous system. Cardiac acceleration is associated with sympathetic activity, whereas deceleration is linked to parasympathetic (vagal) activity. A well-established lateralisation of autonomic processing within the insular cortex holds that the right insula preferentially processes sympathetic cardiac signals, while the left insula preferentially processes parasympathetic signals (Craig 2005; Oppenheimer et al. 1992). Among the electrodes constituting the significant cluster in the present study (FC1, CP1, Pz, CP2, Cz, C4, FC2), right-hemisphere electrodes (C4, FC2, CP2) outnumbered left-hemisphere electrodes (FC1, CP1), indicating a mild right-lateralised topographic distribution. This pattern was consistent with the interpretation that unexpected acceleration — as a sympathetic signal — enhanced HEP amplitude via right insula-associated cortical networks. The relative reduction in HEP amplitude during unexpected deceleration may, in turn, reflect the fact that, under resting-state conditions, sympathetic cardiac signals are processed more strongly within this right-lateralised network than parasympathetic signals.

Compared with unexpected deceleration, unexpected acceleration was associated with stronger beta-band (13–30 Hz) phase synchrony between Cz and Fz, FC2, and F4 during the late time window (250–500 ms; Δwpli = −0.063, p = 0.004). This increase in connectivity was confined to the time window immediately following HEP amplitude modulation (215–250 ms), suggesting that information propagates to a broader cortical network subsequent to primary cardiac signal integration. Beta oscillations have been associated with top-down predictive and regulatory processing (Engel and Fries 2010), and the present finding showed that information exchange between the central (Cz) and frontal regions is facilitated in response to sympathetic signals (unexpected acceleration). The inclusion of right frontal electrodes FC2 and F4 in the frontal cluster is consistent with the aforementioned framework of right insular sympathetic cardiac processing. Taken together, the local HEP amplitude modulation at 215–250 ms and the subsequent increase in right frontal connectivity at 250–500 ms may reflect a sequential cortical elaboration of sympathetic prediction error signals, unfolding from primary cardiac signal integration toward top-down processing mediated by the frontal cortex.

If late frontal processing (250–500 ms) reflects top-down regulatory control, it follows that processing at this stage may be modulated by individual differences in personality. We therefore examined associations between HEP amplitude at the frontal cluster identified in the connectivity analysis (Fz, FC2, F4; 250–500 ms) and personality trait scores. Within this frontal cluster, a significant negative correlation was observed between HEP amplitude during the unexpected acceleration condition and Neuroticism (TIPI_N; ρ = −0.579, p = 0.030): participants with higher Neuroticism scores showed lower frontal cluster HEP amplitude during unexpected acceleration. This association was not observed in the unexpected deceleration condition (ρ = −0.219, p = 0.565), nor was it significant when data were collapsed across all trials — corresponding to conventional HEP amplitude irrespective of prediction error condition (ρ = −0.504, p = 0.099). At least two interpretations of this condition-specific pattern are possible. First, individuals high in Neuroticism may simply have lower cortical sensitivity to sympathetic cardiac prediction errors (unexpected acceleration), which could reflect a neurophysiological trait characterized by attenuated cortical responsiveness to arousal-related cardiac signals. Second, the pattern is amenable to interpretation within the framework of interoceptive avoidance (Khalsa et al. 2018). Interoceptive avoidance refers to a tendency to actively suppress the conscious processing of arousal-related bodily signals, and has been proposed as a transdiagnostic mechanism shared across anxiety spectrum disorders. From this perspective, the selective attenuation of HEP amplitude for sympathetic signals among individuals high in Neuroticism may reflect a motivated inhibitory process that prevents arousal-signaling bodily cues from reaching conscious awareness. Under either interpretation, the present findings suggested that the directional sign of interoceptive prediction error (sympathetic versus parasympathetic) constitutes an important dimension characterizing personality-related individual differences in cortical processing.

As an exploratory finding, a significant negative correlation was observed between HEP amplitude at FC2 (250–500 ms) during trials with large absolute prediction errors and the Future Prediction Task score (FPT score; ρ = −0.455, p = 0.038). Individuals with lower optimism tended to show higher HEP amplitude at FC2 during high-prediction-error trials. Although the interpretation of this finding requires further investigation, it was suggested that cortical sensitivity to the magnitude of cardiac prediction error may be related to individual differences in optimism.

## Acknowledge

This study was supported by grants from the JST Moonshot R&D Grant (JPMJMS2295) and CREST Grant (JPMJCR23P4), JSPS KAKENHI Grant (23H04830, 22K18265, 23K22379), and MEXT Quantum Leap Flagship Program (MEXT QLEAP) Grant (JPMXS0120330644) to MY. JSPS KAKENHI Grant (26K06698) to TH.

## Notes

**Conflicts of Interest:** All authors report no conflict of interest related to this manuscript.

### Competing Interest Statement

The authors have declared no competing interest.

