## Supplemental Figure 1 for "Resting-state heartbeat-evoked potentials are associated with Kalman-derived cardiac prediction errors"

### Supplementary material

#### Results

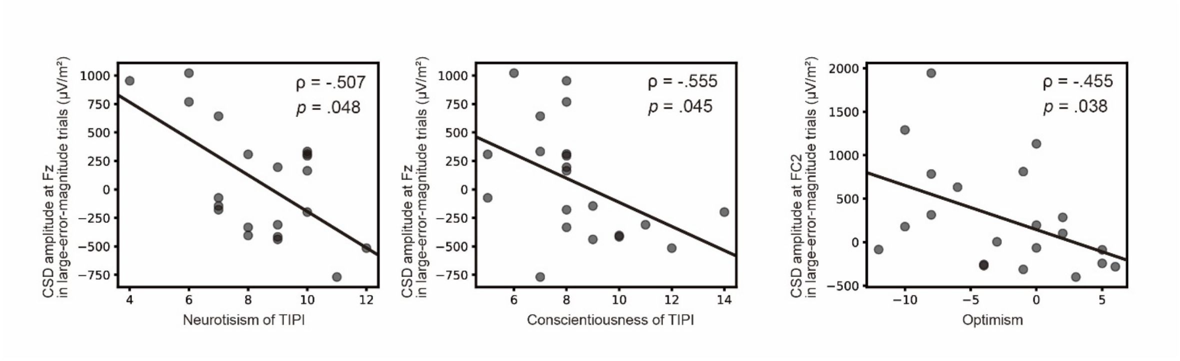


Figure S1. Associations between HEP cluster amplitude in large-prediction-error trials and personality traits. Scatter plots show Spearman correlations between CSD amplitude (μV/m$^2$) at Fz during large-error-magnitude heartbeats and scores on Neuroticism (left) and Conscientiousness (middle), and at FC2 and Optimism (right).
